# The Correlation of Substitution Effects Across Populations and Generations in the Presence of Non-Additive Functional Gene Action

**DOI:** 10.1101/2020.11.03.367227

**Authors:** A. Legarra, C.A. Garcia-Baccino, Y.C.J. Wientjes, Z.G. Vitezica

## Abstract

Allele substitution effects at quantitative trait loci (QTL) are part of the basis of quantitative genetics theory and applications such as association analysis and genomic prediction. In the presence of non-additive functional gene action, substitution effects are not constant across populations. We develop an original approach to model the difference in substitution effects across populations as a first order Taylor series expansion from a “focal” population. This expansion involves the difference in allele frequencies and second-order statistical effects (additive by additive and dominance). The change in allele frequencies is a function of relationships (or genetic distances) across populations. As a result, it is possible to estimate the correlation of substitution effects across two populations using three elements: magnitudes of additive, dominance and additive by additive variances; relationships (Nei’s minimum distances or Fst indexes); and assumed heterozygosities. Similarly, the theory applies as well to distinct generations in a population, in which case the distance across generations is a function of increase of inbreeding. Simulation results confirmed our derivations. Slight biases were observed, depending on the non-additive mechanism and the reference allele. Our derivations are useful to understand and forecast the possibility of prediction across populations and the similarity of GWAS effects.

## INTRODUCTION

One of the aims of quantitative genetics is to provide methods for prediction, for instance genomic prediction (prediction of livestock breeding values or of crop performance) or polygenic risk score (risk of a disease in humans). These predictions would ideally work across a range of populations (different breeds, future generations). Ideally, the prediction goes through a process of identifying causal genes, estimating their effects in some population, and transposing these effects to newly genotyped individuals (Lande and Thompson 1990; Meuwissen *et al*. 2001). These “gene effects” are substitution effects – the regression of the own phenotype (for polygenic risk scores) or expected progeny phenotypes (for estimated breeding values) on gene content at the locus. Being able to use substitution effects at causal genes across populations and generations is a goal of genomic prediction, QTL detection and also of causal mutation finding (Grisart *et al*. 2002).

There are several obstacles for these aims. Finding and validating causal genes and understanding their functional mechanism is extremely difficult (Grobet *et al*. 1997; Bonifati *et al*. 2003; Rupp *et al*. 2015). In practice, predictions are done using SNP markers using statistical genetics techniques. In livestock, use of markers results in very good predictions within populations, but mediocre (at best) predictions across populations, even with very sophisticated techniques (Hayes *et al*. 2009; Karoui *et al*. 2012; Porto-Neto *et al*. 2015; MacLeod *et al*. 2016). Indeed, livestock and human genetics empirical results show decreasing predictive ability with increasing genetic distance across distinct populations or generations (Liu *et al*. 2016; Martin *et al*. 2019). In humans, there is strong correlation between GWAS effect estimates across human populations, with typical values around 0.82 (Marigorta and Navarro 2013; Shi *et al*. 2021). The lack of perfect linkage disequilibrium (LD) across markers and genes has been claimed to be a reason for this decrease in accuracy. Adding extra information (more dense maps, biological prior information) should result in a better choice of markers close to causal genes, and therefore in a boost in predictive abilities across populations (Roos *et al*. 2009; MacLeod *et al*. 2016). However, in practice, the increase in predictive ability across populations is small at best (MacLeod *et al*. 2016; Moghaddar *et al*. 2019). This is against results from simulations (Roos *et al*. 2009; MacLeod *et al*. 2016) – but a problem in simulations is that typically gene effects are assumed to be biologically additive and therefore constant across populations.

We argue that, although imperfect LD is a likely cause for not being able to predict across populations, it is not the only one. In fact, substitution (statistical) additive gene action is not homogeneous across populations, even for exactly the same causal mutation. Examples in livestock genetics include myostatin gene (Aiello *et al*. 2018) or DGAT1 (Gautier *et al*. 2007). For instance, in the latter, the “K” allele had rather different substitution effects across breeds: −611, −142 and −351 kg of milk for the respective breeds Montbéliarde, Normande and Holstein, for a trait with a genetic standard deviation of ~600 kg. Although part of these differences may be due to genotype by environment interactions, it is also plausible that this is due to epistasis or dominance; for instance this is the case in DGAT1 (Streit *et al*. 2011). For instance, in the 5-loci epistatic network in Carlborg et al. (2006) some substitution effects at genes switch signs depending on genetic background. In Drosophila, estimated substitution effects of P-element insertions switched signs depending on the genetic background (Magwire *et al*. 2010; Mackay 2015).

There is indeed widespread evidence of biological epistasis (Mackay 2014). Whereas biological epistasis does not impede (on the contrary) large additive variation (Hill *et al*. 2008; Mackay 2014; Mäki-Tanila and Hill 2014), it does imply that substitution effects do vary across genetic backgrounds. Thus, in the presence of functional dominance and epistasis, there is no stability of substitution effects across different genetic backgrounds. Even if in all of these populations, additive variation accounts for most genetic variation, and additive substitution effects are sizable (Hill *et al*. 2008), substitution effects may differ across populations.

Recent simulations (Dai *et al*. 2020; Duenk *et al*. 2020) showed that the difference in substitution effects across populations may be quite large under non-additive biological gene action and increases with divergence of populations. It is relatively easy to derive algebraic expressions for substitution effects *α* assuming specific hypothesis of biological gene action. For instance, assuming biological additive and dominance effects only (Falconer and Mackay 1996) results in *α* = *a* + (*q* – *p*)*d*, whereas assuming additive, dominance and additive by additive biological gene effects results, assuming linkage equilibrium (LE), in *α*_1_ = *a*_1_ + (*q*_1_ – *p*_1_)*d*_1_ + (*p*_2_ – *q*_2_)[*aa*]_12_ (Fuerst *et al*. 1997). Both of the approaches (simulation or analytical) are limited because the hypotheses of specific biological gene actions *(e.g*. 2-loci interaction but not 3-loci interaction) are too restrictive. Classically, these hypotheses are bypassed in quantitative genetics by working on statistical effects (Falconer and Mackay 1996; Lynch and Walsh 1998; Mäki-Tanila and Hill 2014).

The key parameter to describe the resemblance of statistical effects across populations is the correlation of substitution effects across populations. Under certain assumptions (independence of allele frequencies and substitution effects, appropriate coding; (Wientjes *et al*. 2017)), this correlation can be estimated from SNP markers and data of two populations (Karoui *et al*. 2012; Wientjes *et al*. 2017), and can easily be accommodated into genomic prediction models (Karoui *et al*. 2012; Xiang *et al*. 2017). In human studies, the correlation is estimated through the meta-analysis of GWAS statistics (Marigorta and Navarro 2013; Shi *et al*. 2021). However, in addition to empirical results, some theory to describe the resemblance of substitution effects across different populations would be helpful to (1) better understand and quantify the change in *true* substitution effects *(i.e*. if true genes instead of markers were being used), (2) give upper bounds for genomic prediction across populations/generations, and (3) allow *a priori* planning of genomic predictions *i.e*. to include or not different subpopulations.

The aim of this work is to develop a theory to understand and predict, under a neutral scenario, the extent of change of substitution effects across space (breeds, lines) and across time (generations), without invoking or assuming specific modes of biological gene action. As a result, we obtain explicit estimators that are functions of additive, dominance and additive by additive variances, genetic distances across populations, and distributions of allele frequencies. We check and illustrate our theory using published results and simulations considering dominance and epistasis (additive by additive and complementary) from 5-loci interactions. Main factors affecting correlation of substitution effects across populations are their genetic distance and the extent of additive by additive variation, which is rarely large.

## THEORY

### Analytical results

#### General Theory

Here, we model difference of substitution effects across two populations as Taylor expansions around one of them, the “focal” population. Using Kojima’s method (Kojima 1959, 1961), we put additive substitution effects as a function of differences in allele frequencies across populations, “focal” additive substitution effects, and second order (dominance, additive by additive epistasis) statistical effects. From here, we show that the correlation of substitution effects across populations is (approximately) a function of their differentiation (or genetic distance), the additive, dominant and additive by additive genetic variances, the average heterozygosities and average squared heterozygosities. All these parameters can be estimated in real populations. In the following, we try to stick to Mäki-Tanila and Hill (2014) notation. Many details are given as Appendix in the Supplementary Material. Main notation is presented in Table 1.

**Table 1.** Notation.

|  |  |  |  |
| --- | --- | --- | --- |
| $\alpha_i^b, \alpha_i^{b'}$ | Substitution effect of locus $i$ in population $b$, in population $b'$ | $\sigma_{\alpha,b}^2$ | Variance of substitution effects in population $b$ |
| $d_i^{*b}$ | Dominance deviation at locus $i$ in population $b$ | $\sigma_{d,b}^2, \mu_{d,b}$ | Variance and mean of dominance deviations in population $b$ |
| $(\alpha\alpha)_{ij}^b$ | Additive by additive effect of loci $i, j$ at population $b$ | $\sigma_{(\alpha\alpha,b)}^2$ | Variance of additive by additive effects in population $b$ |
| $(\alpha\alpha)_i^b$ | Vector of additive by additive effects of locus $i$ with all other loci in population $b$ | | |
| $p_i^b, p_i^{b'}$ | Allele frequency of locus $i$ in population $b$, in population $b'$ | | |
| $\mathbf{p}^b, \mathbf{p}^{b'}$ | Vectors of allele frequencies in population $b$, in population $b'$ | | |
| $\bar{H}_b$ | Average heterozygosity at population $b$, $\bar{H}_b = E(2p_i^b(1 - p_i^b))$ | | |
| $\bar{H}_b^2$ | Average squared heterozygosity at population $b$,<br>$\bar{H}_b^2 = E\left(\left(2p_i^b(1 - p_i^b)\right)^2\right)$ | | |
| $\overline{HH}_b$ | Average product of heterozygosities at population $b$,<br>$\overline{HH}_b = E_{i>j}\left(4p_i^b(1 - p_i^b)p_j^b(1 - p_j^b)\right)$ | | |
| $\epsilon_i$ | Difference in allele frequencies at locus $i$,<br>$\epsilon_i = p_i^{b'} - p_i^b$ | $\sigma_\epsilon^2$ | Variance of the difference in allele frequencies across all loci |
| $D_{b,b'}$ | Nei's minimum genetic distance | | |
| $\sigma_A^2, \sigma_D^2, \sigma_{AA}^2$ | Additive, Dominance and Additive by Additive variances (in population $b$) | | |

Note that our procedure is general – it does not invoke any particular mechanism for epistasis or dominance, nor knowledge of individual QTL effects and locations. We assume that the population mean is a (possibly complex) function of QTL allele frequencies ***p*** and QTL functional or biological effects. The latter, albeit unknown, are assumed to be constant across populations – we therefore do not consider genotype by environment interactions.

Consider the correlation 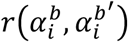 of substitution effects *α^b^* and *α^b′^* across respective populations (breeds, heterotic groups, lines, or generations) *b* and *b′*. For simplicity, the allele to which *α* refers is random – it can be either the wild or the mutant allele. In this manner the average value of *α* is 0, even in presence of deleterious mutations. It is known that, in presence of dominant and epistatic biological interactions, the value of *α* depends on the allele frequencies and biological effects; for instance, *α*_1_ = *a*_1_ + (*q*_1_ – *p*_1_)*d*_1_ + (*p*_2_ – *q*_2_)[*aa*]_12_ for 2-loci epistasis (Fuerst *et al*. 1997), for *p*_1_ = 1 – *q*_1_ and *p*_2_ = 1 – *q*_2_ frequencies at loci 1 and 2, respectively. Inspired by this, we want to write the correlation 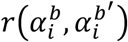 for a locus *i* as a function of vectors of respective allele frequencies, ***p***^*b*^ and ***p***^*b′*^, but in a general manner, without defining a particular functional or biological gene action. We use a first order Taylor series expansion to approximate additive substitution effects in a population *b′*, as a function of effects in another “focal” population *b* and their distance. By doing this, it can be shown that the difference of substitution effects between two populations is (approximately) a function of the genetic distance of the two populations, and the magnitude of dominance and second order epistatic variances in the focal population *b*.

To derive additive substitution effects *α* as function of allelic frequencies ***p***, we use Kojima’s definition of statistical effects as first, second… derivatives of the mean of the population (*μ*) as a function of *p*. We don’t invoke any explicit function – we just presume that there is one, in other words, change in allele frequencies of the population implies change in the total average genotypic value. Using Kojima’s method, the additive substitution effect at the *i*-th locus is the first derivative (Kojima 1959, 1961):

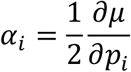

Higher order statistical effects implying locus *i* (i.e. dominance deviations and epistatic interactions) can be represented by higher order partial derivatives of *μ* or equivalently as derivatives of *α_i_*. The dominance deviation at the *i*-th locus (that we denote as *d*\* to distinguish from the biological or functional effect *d* (Falconer and Mackay 1996)) is:

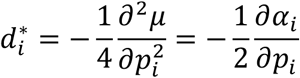

The negative sign comes because the dominance deviation is usually understood as a feature of *heterozygosity*, in other words, it is of opposite sign than the increase of homozygosity in 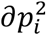. Last, the epistatic pairwise deviation of locus *i* with *j* is

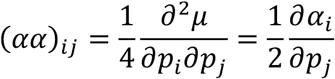

This is positive because it is the effect of increasing both *p_i_* and *p_j_*. Note that interaction of order *k* implies k-ith order derivative with scaling factor 1/2^*k*^.

Kojima’s method shows, therefore, that higher order effects of one locus are derivatives of lower order effects. With these elements we can make a Taylor order expansion of *α_i_* around frequencies in the “focal” population, ***p**^b^*, so that ***p**^b′^* = ***p**^b^* + ***ϵ***, such that from values of *α* in *b* and changes in allele frequencies ***ϵ*** = ***p**^b′^* – ***p**^b^* we create a function 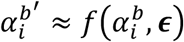. In the Appendix (section 1.1), we show that the Taylor linear approximation of the substitution effects of population *b*′: 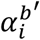, from effects from populations 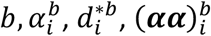 is:

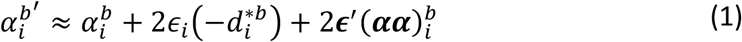

where we use differences in allele frequencies ***ϵ***, 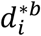 is the statistical dominance deviation at the locus *i* and 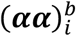 is a vector containing epistatic substitution effects of locus *i* with the rest of loci. By convention we assign 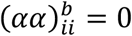.

From equation 1, the covariance across two populations *b* and *b′* of the two substitution effects 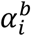 and 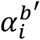 of the locus *i* is:

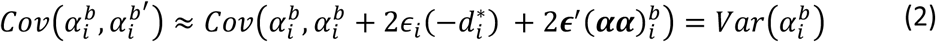

The equality holds because terms 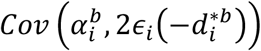 and 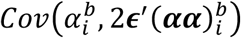 are null, given that the different statistical effects (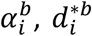 and 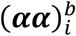) are mutually orthogonal by construction.

Thus 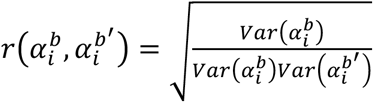, and now we need the variance of 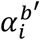 as a function of effects in population *b*, this is (see the Appendix, section 1.2 for details):

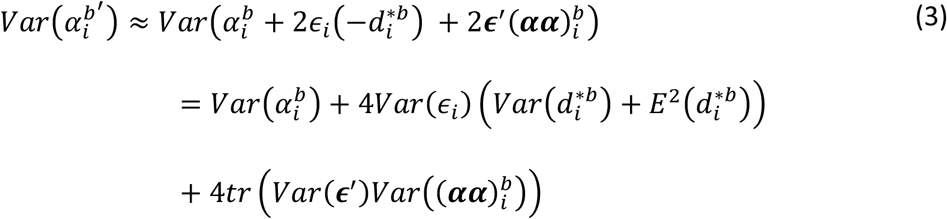

This expression is unsymmetric and seems to imply that by construction 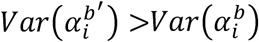; the reason for this is that for the “focal” population *b* the variance (or at least its estimators) is better known than for the “approximated” population *b′*. An alternative derivation in the Appendix (section 1.4) shows that, if the “focal” population is a third one (*f*) a set of expressions analogous to (2) and (3) is:

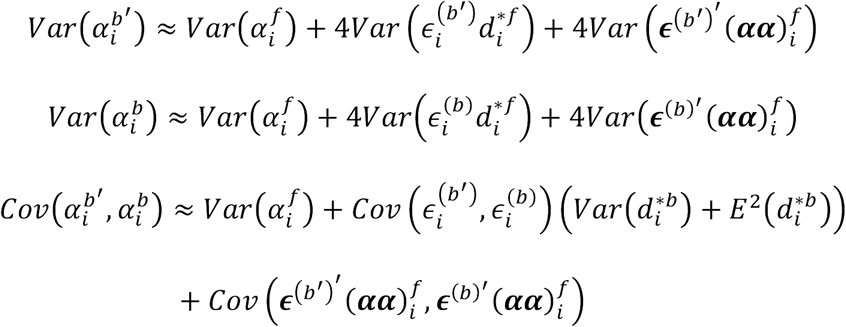

Where 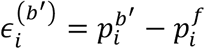 and 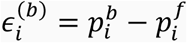. From these expressions, we obtain (2) and (3) as a particular case if the focal population is *b*.

In the expression (3), *Var*(***ϵ***′) and 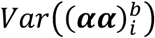 are matrices. The first one describes the variability of differences in allele frequencies:

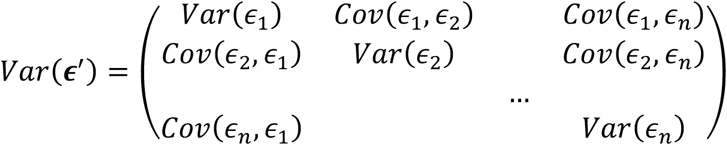

A locus that may diverge a lot (for instance because it is highly polymorphic) has high *Var*(*ϵ*); two loci in strong linkage will show non-zero *Cov*(*ϵ*_1_, *ϵ*_2_). The second matrix contains (co)variances of the epistatic effects across all pairs marker *i* – other markers:

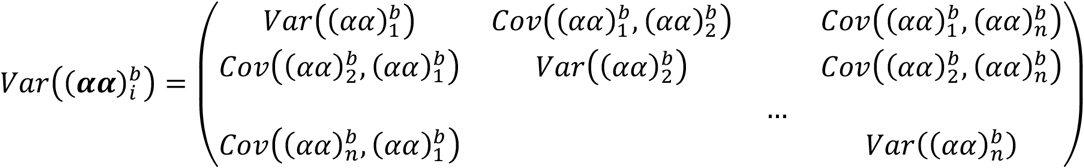

For instance, 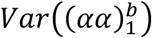 contains the variance of the epistatic effect of marker i with marker 1, and so on. We assume 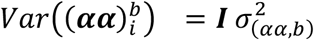, *i.e*. epistatic terms 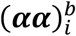 can be either positive or negative, with a variance 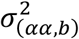 and are a priori uncorrelated to each other. Assuming null off-diagonals for 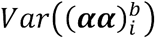 results that, in the previous product 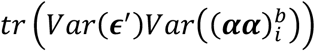, off-diagonal elements of *Var*(***ϵ***′) disappear from the result (even if they are not null). Thus, assuming that all diagonal elements of *Var*(***ϵ***′) have the same common variance 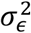, this results in 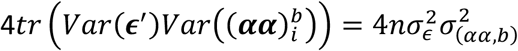. Note that assuming that all diagonal elements of *Var*(***ϵ***′) are equal is an approximation – for instance more polymorphic loci vary more.

The expression 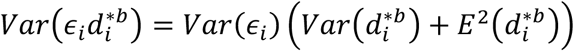 is detailed in the Appendix (section 1.2), and it shows that both the variability of dominance deviations 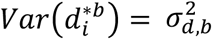, and its mean 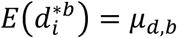 which, if not zero, can be understood as the basis of inbreeding depression, enter into the expression. Also, we assume ***ϵ*** (change in allele frequencies, but not the allele frequencies themselves) and the different non-additive effects 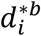 and 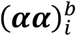 to be independent (uncorrelated). This makes sense as loci may be selected for additive effects but not for non-additive effects.

Our next goal is to relate these results in (3) to quantities that are measured empirically. In particular, we need the variances of the different genetic effects and the variance of changes in allele frequencies. We address these two terms in turn.

We factorize the variance of statistical additive, dominant and additive by additive effects as follows (Mäki-Tanila and Hill 2014; Vitezica *et al*. 2017):

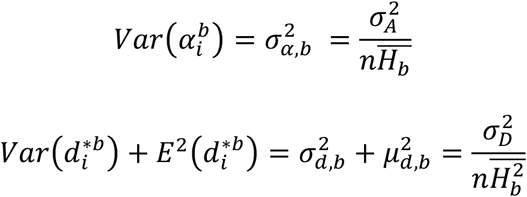

(the latter is shown in the Appendix, section 1.5) and

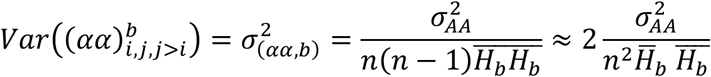

for *n* the number of QTL loci and using functions of heterozygosities (more details in the Appendix, section 1.6):

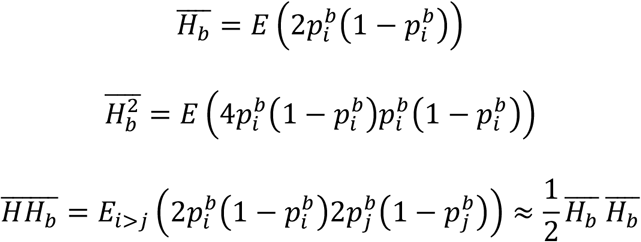

Here we have assumed independence of QTL allele frequencies and QTL effects. All variances and effects, as well as heterozygosities *H_b_*, refer to the focal population *b* with allele frequencies *p^b^* and effects *α^b^*. Note that we assume HWE and LE within both *b* and *b*’. Of course, we don’t know allele frequencies or even numbers of true causal genes, but the first can be prudently guessed and the latter cancel out in the following.

Consider the scalar 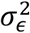, the variance of differences of allele frequency. In fact, 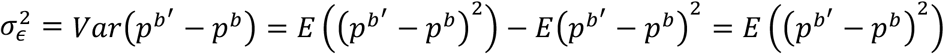 because 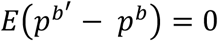 when averaged across loci. Therefore, 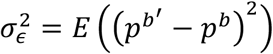 which corresponds to Nei’s “minimum genetic distance” (Nei 1987; Caballero and Toro 2002), and, accordingly, will be called 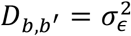. The value of *D_b,b′_* can be estimated from marker data as 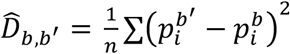, although it is sensitive to the spectra of polymorphisms (e.g. SNP chips vs sequencing). In addition, *D_b,b′_* is also the numerator of the *F_ST_* fixation index, e.g. Hudson’s(1992) 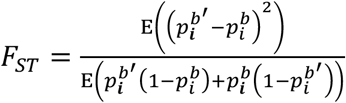 with an estimator 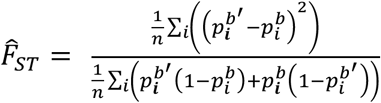 (Hudson *et al*. 1992; Bhatia *et al*. 2013). In principle, *D_b,b′_* and *F_ST_* can be estimated from markers, but also from evolutionary distances and effective population size, or both (Weir and Hill 2002; Bonhomme *et al*. 2010). Now we can rewrite the last term of equation (3) as

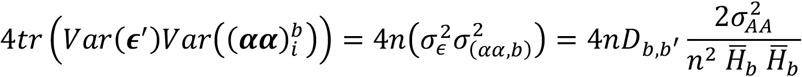

Combining expressions above, equation (3) becomes

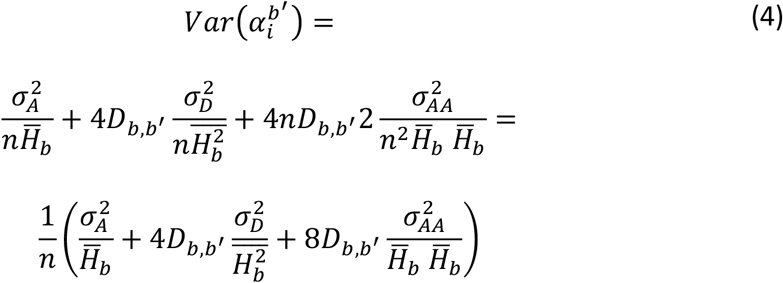

From here the correlation of *α* across populations is (factorizing out 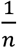)

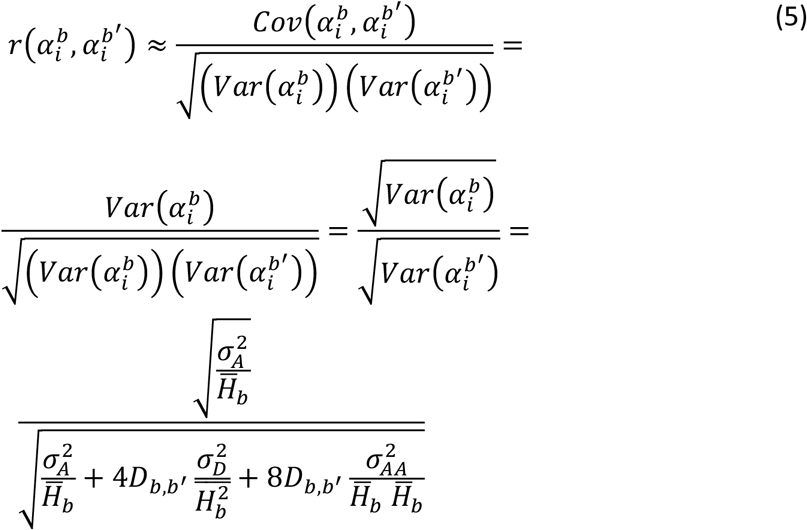

Expression [5] can also be obtained if we start the Taylor expansion from a third focal population that is neither *b* nor *b′*, as shown in the Appendix, section 1.4. Factorizing out 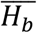 we arrive to the slightly clearer expression

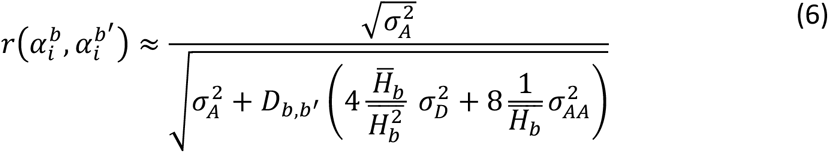

which shows well that the correlation is a function of distance across populations (*D_b,b′_*) and weights of additive *vs*. non-additive variances. Note that *D_b,b′_* is always positive, which implies that 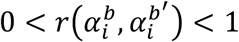 as expected.

The quantities involved in equation (6) are (1) Nei’s “minimum genetic distance” *D_b,b′_*, which describes the similarity of populations *b* and *b′*, (2) the variance of statistical additive, dominant and additive by additive effects at the individual level 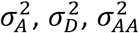 in population *b* (3) first and second moments of heterozygosities 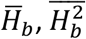. All these values can be, in principle, estimated from data or “prudently guessed”. For the particular case of heterozygosities, SNP markers are a poor choice and it is likely better to use a guess based on sequence or evolutionary processes (coalescence).

From the definition of *F_ST_* and assuming 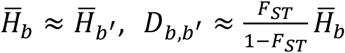 (detailed in the Appendix, section 1.7), leading to

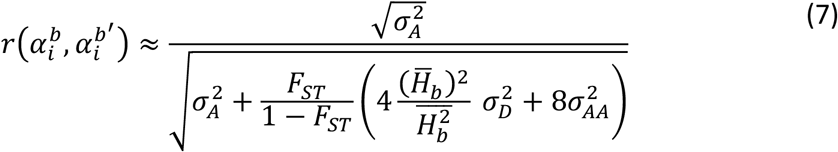

The advantage of the *F_ST_* is that it is more robust to the spectra of allele frequencies used to estimate it (Bhatia *et al*. 2013). If we further assume that dominance variance 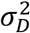 is negligible, we can write a neat expression of the correlation in terms of *F_ST_*:

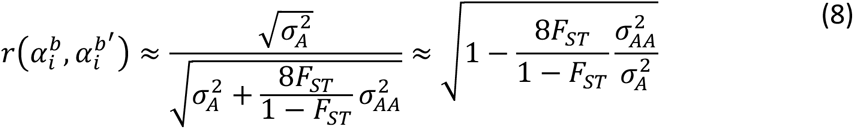

The second approximation involving 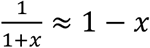. This expression (8) plainly tells that the squared correlation of gene substitution effects across two population is (to a few degrees of approximation) a linear function of the similarity of populations and the additive by additive variance. To our knowledge, these results showing the importance of the different factors on the difference between substitution effects had been shown before only through simulations. Thus, the algorithm to estimate *a priori* the correlation of *α* across populations *b* and *b’*, 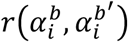, is:

1. Estimate in population *b*

a. additive, dominance, and additive by additive variances
b. average heterozygosity 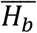 and average squared heterozygosity 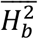
2. Estimate Nei’s distance *D_b,b′_* and/or *F_ST_* of the two populations
3. Apply equation (6) or (7).

#### Consideration of directionality of substitution effects

In Plant and Animal Breeding the origin of the allele is often overlooked, as a mutation may be evolutionary harmful but of interest for farming, and also because many traits selected for do not have a close relationship to fitness in the wild. However, in Evolutionary Genetics it is reasonable to think that most mutations are deleterious, thus with a negative effect of the mutant allele. In Medical Genetics reports of estimated substitution effects are also often done in terms of “susceptible” alleles. In both cases *E*(*α*) ≠ 0 or even *α* > 0 for all loci. In both cases *α* is “oriented” and has no zero mean. In the theory above we have considered that *α* is the effect of a randomly drawn allele, which leads to *E*(*α*) = 0 and enormously simplifies the algebra. In order to consider oriented *α*, we propose to transform the estimate of 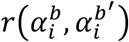 into 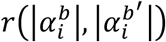, the correlation of the absolute values. Assuming that *α* follows a normal distribution, |*α*| follows a so-called folded normal distribution. From here, 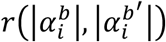 is obtained from 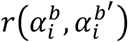 using expressions (not detailed here) in Kan and Robotti (2017), conveniently programmed in the R package MomTrunc (Galarza *et al*. 2020). The specific R function is in the Appendix, section 1.8, and we will call it r2rabs().

#### Covariance across generations within one population

The definition above of two populations is general enough that we can consider any two populations, *e.g*. two breeds (Angus and Hereford), two strains (New Zealand Holstein and US Holstein) or two generations or time frames (*e.g*. animals born in 2000 vs. animals born in 2005, or animals born in 2000 vs. their descendants). There is evidence that across-generations genetic correlation decreases with (many) generations to values as low as 0.6 (Tsuruta *et al*. 2004; Haile-Mariam and Pryce 2015). Part of this is likely due to genotype by environment interactions. Anyway, part of the across-generation genetic correlation could be due to changes in the allele frequency due to drift, and therefore it can be accounted for by our model, based on the evolution of average coancestry in the breed. We develop this next.

We will talk about “generations” but ideas apply to pedigrees with overlapping generations as well. Consider that what we previously called populations *b* and *b′* are animals born at time *t*_1_ and *t*_2_, with *t*_2_ > *t*_1_. Equation (7) becomes 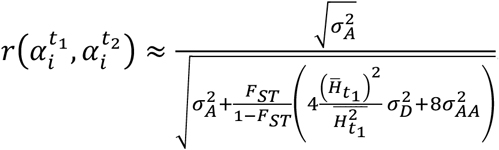. In pedigree-based context, the *F_ST_* is simply half the increase in average relationship from *t*_1_ to *t*_2_ (Powell *et al*. 2010) which is approximately equal to the increase in inbreeding from *t*_1_ + 1 to *t*_2_ + 1, which in turn is approximately (*t*_2_ – *t*_1_)Δ*F* for small values of Δ*F* and steady increase of inbreeding. Thus, 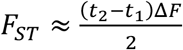 to obtain

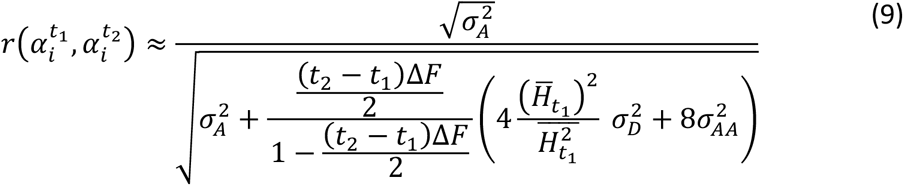

Assuming, like in (8), small values of dominance variance and of *F_ST_*, we obtain

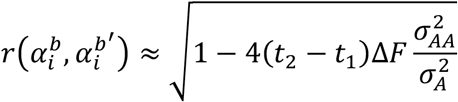

Thus, the correlation of substitution effects decreases when there is large drift, reflected in high values of Δ*F*. This is the case for instance if parents of the next generation are very highly selected without restrictions in future inbreeding, resulting in a considerable change in allele frequencies over the generations.

This agrees with classical theory (Falconer and Mackay 1996): the variance in change of allele frequencies from one generation to the next is simply Δ*F*. Typical values of Δ*F* in livestock (rabbits, pigs, cattle or sheep) are at most 0.01 per generation (Welsh *et al*. 2010; García-Ruiz *et al*. 2016; Fernández *et al*. 2017; Rodríguez-Ramilo *et al*. 2019), although this is potentially changing with genomic selection.

### Simulations

#### Description of the simulations

The objective of the simulation is to verify our results considering different kinds of non-additive biological gene action, and not to infer the values of the correlation of substitution effects across populations as this requires realistic presumptions of the forms and magnitudes of epistatic interactions, something that is largely unknown.

We used *macs* (Chen *et al*. 2009) to simulate a “cattle” scenario of two domestic cattle breeds which diverged from a common ancestral population, in the lines of (Pérez-Enciso 2014; Pérez-Enciso *et al*. 2015), where a large population had a 10-fold reduction bottleneck (domestication) 2000 generations ago and a split into two populations *t* generations ago, where we made *t* oscillate between 0 and 100 by steps of 10. Parameters in the simulation were tailored (Pérez-Enciso 2014; Pérez-Enciso *et al*. 2015) to mimic observed levels of diversity in cattle (Gibbs *et al*. 2009) and lead to *F_ST_* values between 0 and 0.15. We considered 100 DNA stretches of 300 Kb each. We simulated 200 individuals per population, from which we obtained allele frequencies per population. Details are provided in the Appendix, section 1.9. This provided ~400,000 segregating loci with realistic allele frequencies (L-shaped distribution of allele frequencies).

To simulate non-additive biological gene action, we considered three scenarios. All of them involved 5000 single marker polymorphisms drawn at random from the simulated ones with no particular restriction. First scenario was Complete Dominance (within locus); second one was 1000 networks of 5 loci in Complementary Epistasis, and the third one 1000 networks of 5 loci with Multiplicative Epistasis (additive by additive). We also simulated networks of 2 and of 10 loci with similar results (not shown). These scenarios are similar to Hill *et al*. (2008) but instead of considering 2 loci we consider 5. We use these forms of epistasis, as the first two ones may be interpreted as biologically meaningful gene actions, and the third one is an extreme case for the change in substitution effect across populations. In addition, all three scenarios are analytically tractable. Complete dominance has the genotypic value of the heterozygote equal to one of the homozygotes, e.g. the presence of a single copy of the “good” allele is enough to, say, avoid the disease. Complementary epistasis can be seen as a multi-loci dominance, e.g. disease happens when there is a recessive deleterious genotype at *any* of the loci. Additive by additive epistasis can be understood as a pure multiplication of gene contents and has no good biological interpretation. Networks contribute additively to the total phenotype. Note that, by definition, functional gene action is the same across all populations. From the description of the non-additive gene action and some algebra in the Appendix, we are able to derive analytically the values of *α, d*\* and (*αα*) in each population and variances 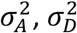 and 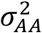. Details are given in the Appendix, section 1.10. For each value of *t*, we made 10 replicates and averaged the results.

From the true substitution effects *α^b^* and *α^b′^* derived above, we obtained the true value *r*(*α^b^, α^b′^*). Note that because of the coalescent simulation, all reference alleles (i.e. with frequency *p*) are “mutant” ones, and due to assumed dominance and epistatic actions, *α* are negative by construction (i.e. the mutant allele is deleterious). To conciliate this fact with our derivations, that assume that *α* refers to a random allele and has null means, we did two things: (1) compute a “random allele” version of *α^b^, α^b′^* in which *α* was changed sign for “odd” loci, and (2) estimate *r*(*α^b^, α^b′^*) using the transformation of normal distribution into folded normal (Kan and Robotti 2017), i.e. the r2rabs() function mentioned above. Thus, we have two estimands, the correlation for the “mutant allele” effect *r_mutant_*(*α^b^, α^b′^*) (which corresponds *e.g*. to typical use in Evolutionary and Medical Genetics) and the correlation for the “random allele” effect *r_random_*(*α^b^, α^b′^*) (which corresponds, *e.g*. to genomic selection and some GWAS). We observed that *r_mutant_*(*α^b^, α^b′^*) < *r_random_*(*α^b^, α^b′^*). For instance, in one of the scenarios *r_mutant_*(*α^b^, α^b′^*) = 0.67 and *r_random_*(*α^b^, α^b′^*) = 0.85.

Now we describe the estimators. We considered either the use of “all” polymorphisms, or of a “SNP” selection in which MAF>0.01 across both populations simultaneously (roughly 40,000 polymorphisms), as this corresponds to typical SNP panels in genetic improvement. Then we used the equation (6) either for “all” or for “SNP”, using their frequencies to obtain *D_b,b′_*, 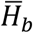 and 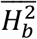. In “all”, the spectra of allele frequencies of polymorphisms and of QTL is the same, but not in “SNP”. As “SNP” tends to be biased, we also considered the equation (7) using *F_ST_* estimated from SNPs and 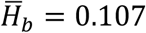, and 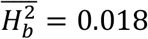 from the “Hill” U-shaped distribution mentioned above. Note that *F_ST_* is more robust to the spectra of allele frequencies used to estimate it. Thus, we obtained three estimators for “random allele” 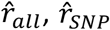 and 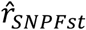, and also the three corresponding estimators for “mutant allele” *r_mutant_*(*α^b^, α^b′^*) applying function r2rabs() to 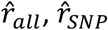 and 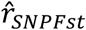.

#### Results of the simulations

Simulated variance components (expressed as ratios from total genetic variance *σ*^2^) with marker genotypes and with QTL genotypes are shown in Table 2. All scenarios yield high additive genetic variances, as expected.

**Table 2.**
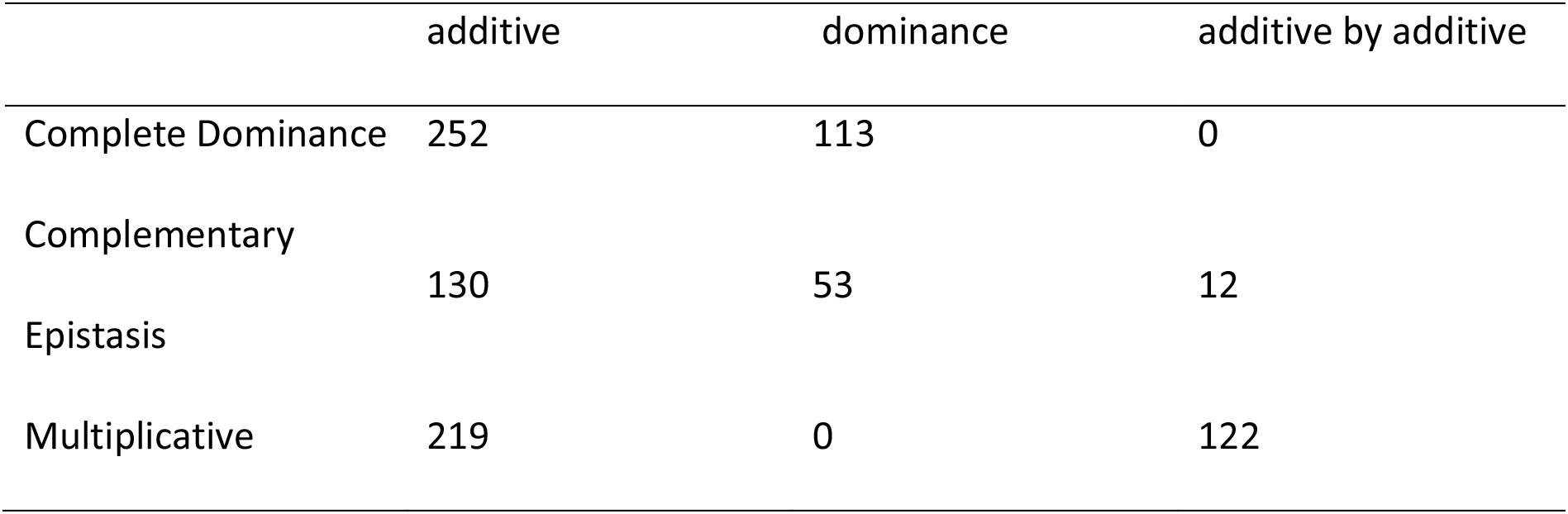
Additive, dominance and additive by additive variances in the simulated population

True simulated values of correlation across substitution effects *r_random_*(*α^b^, α^b′^*) and their estimates 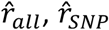 and 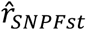, are presented in Figure 1 (for *α* defined for random alleles) and Figure 2 (for *α* defined for mutant alleles). Generally speaking, Figure 1 applies to nonfitness related traits (for “random”) and Figure 2 to fitness-related traits (for “mutant”). As predicted by our derivations, there is a clear and almost linear decrease of *r*(*α^b^, α^b′^*) with increasing values of *F_ST_*.

**Figure 1.**
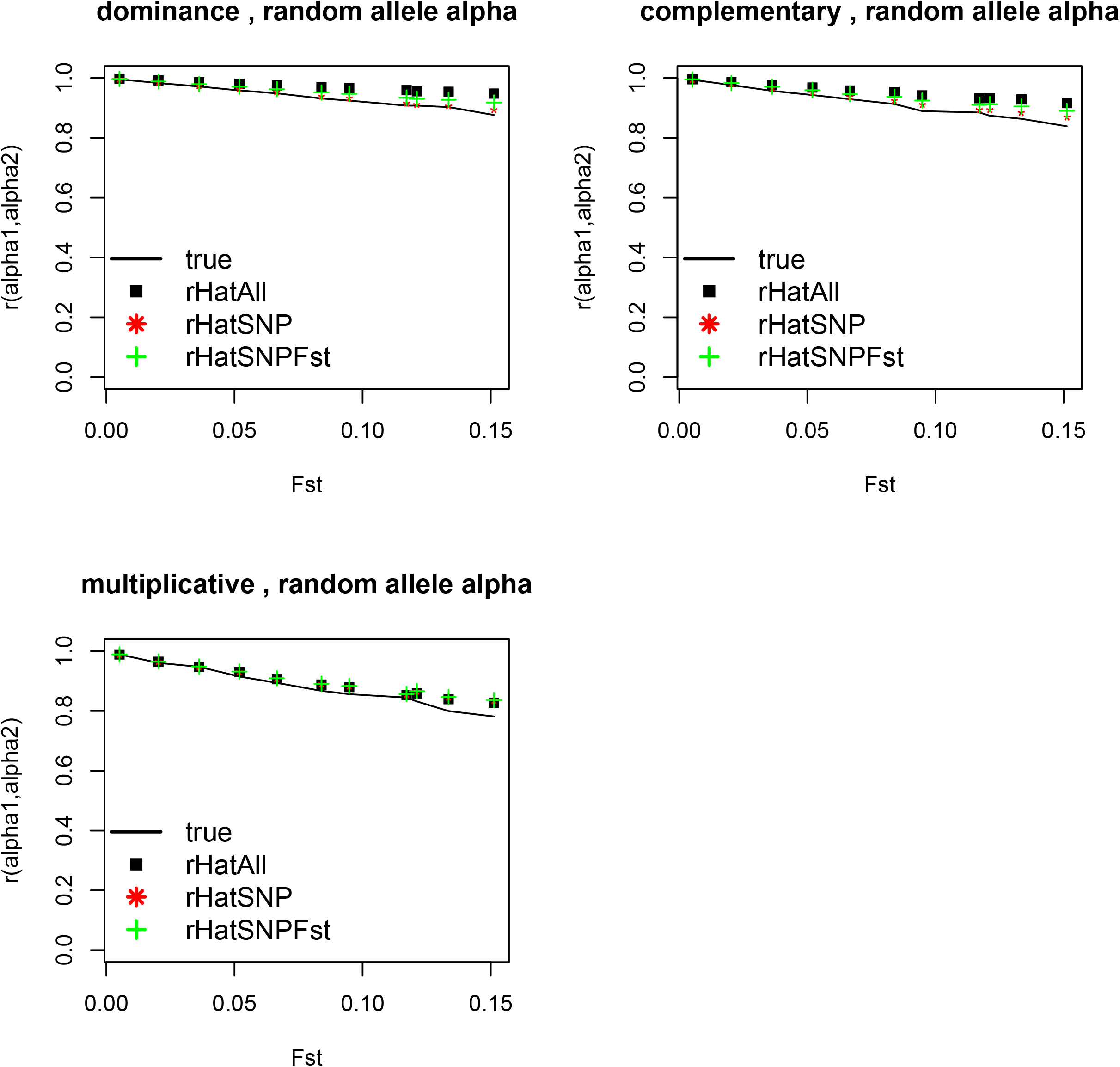
Simulated (straight black line) and estimated (points) correlation across QTL substitution effects of random alleles across two populations as a function of their *F_ST_* differentiation coefficient. rHatAll: estimates with all polymorphism. rHatSNP: estimates using SNP-like loci. rHatSNPFst: estimates using SNP-like loci with a correction for heterozygosity. Results of 10 replicates per point with s.e.<0.01.

In Figure 1 (for *α* defined for random alleles) it can be observed that our expressions (6–7) tend to slightly over-estimate *r_random_*(*α^b^, α^b′^*) for Complete Dominance, and Complementary Epistasis. The estimates are quite good for Multiplicative Epistasis. Estimators using SNP markers are slightly less accurate than using the complete polymorphism spectra, and using *F_ST_* from SNP markers, coupled with a guess of heterozygosities, is similar to using SNP alone to infer both distances and heterozygosities.

Figure 2 (for *α* defined for mutant alleles) presents values of *r_mutant_*(*α^b^, α^b′^*) and estimates 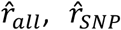 and 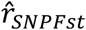. Estimating *r_mutant_*(*α^b^, α^b′^*) is more difficult than estimating *r_random_*(*α^b^, α^b′^*): depending on the scenario there is more over- or under-estimation that for *r_random_*(*α^b^, α^b′^*). In this case the effect of using all polymorphisms or SNP is more marked. However, overall, we find that our estimators explain well the decay in *r*(*α^b^, α^b′^*). The imperfect disagreement (both in Figures 1 and 2) is probably due to several wrong assumptions; we comment some of them in the Discussion.

**Figure 2.**
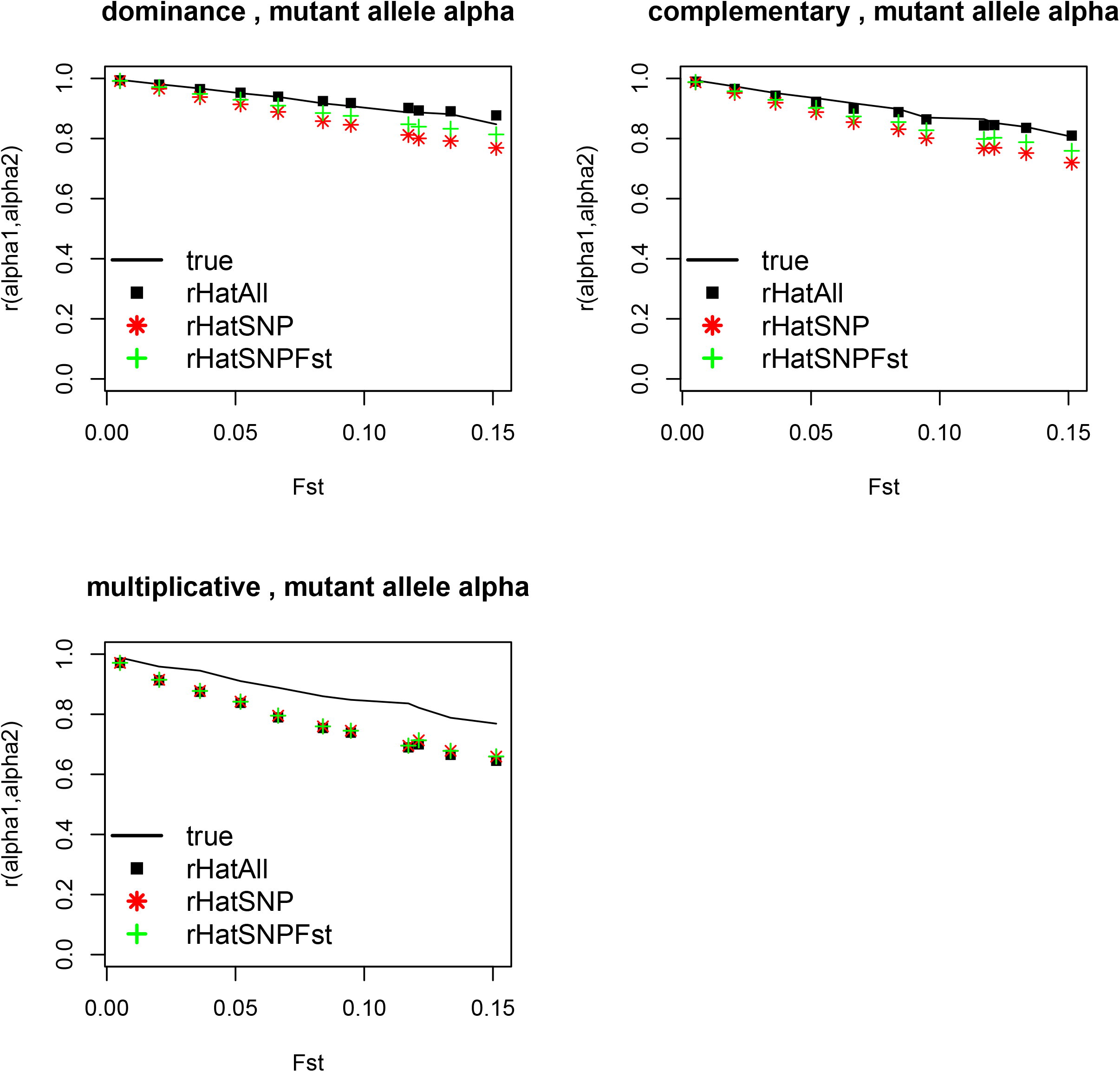
Simulated (straight black line) and estimated (points) correlation across QTL substitution effects of mutant alleles across two populations as a function of their *F_ST_* differentiation coefficient. rHatAll: estimates with all polymorphism. rHatSNP: estimates using SNP-like loci. rHatSNPFst: estimates using SNP-like loci with a correction for heterozygosity. Results of 10 replicates per point with s.e.<0.01.

## EMPIRICAL EXAMPLES

### Estimates of correlation across populations using literature values

#### Literature values used and assumptions

From values of the literature, the previous expressions, equations (6) and (7) were applied to obtain guesses of the correlation of substitution effects across different populations and generations within population. Estimates of statistical non-additive variation are scarce, as are estimates of distances across populations in terms of covariance of allele frequencies. The differentiation *F_ST_* of Jersey and Holstein breeds of cattle obtained from (VanRaden *et al*. 2011) is 0.06, and among Landrace and Yorkshire is 0.16 (Xiang *et al*. 2017).

Table 3 includes literature estimates for additive, dominance and epistatic variation. We used estimates for milk yield in Simmental cattle (Fuerst and Sölkner 1994) as if it was Holstein, and for litter size in a commercial pig line (Vitezica *et al*. 2018) as if it was Landrace, because we could not find estimates in the same populations. These estimates (as most estimates of nonadditive variances) are inaccurate and are presented here only as examples. In the particular case of cattle, the estimated value of 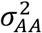 is higher than usually expected *a priori* (Hill *et al*. 2008), so we take it as an example of an extreme, but not impossible, case.

**Table 3.** Variance component estimates, as ratio from phenotypic variance, from literature

| Species | $\sigma_A^2$ | $\sigma_D^2$ | $\sigma_{AA}^2$ |
| --- | --- | --- | --- |
| Cattle | 0.20 | 0.09 | 0.15 |
| Pigs | 0.092 | 0.020 | 0.016 |

Then, we assumed three distributions for QTL allele frequencies, following Hill *et al*. (2008): a uniform distribution (“Uniform”), and a U-shaped distributions with *f*(*p*) ∝ *p*^−1^(1 – *p*)^−1^ with effective population size of *Ne* = 50 (“Hill”), bounded at 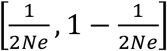. We also assumed a Beta(0.04, 0.04) distribution, which is an extreme U-shaped distribution (“Extreme”) in which roughly 80% of the QTLs have minor allele frequency lower than 0.01. This results in respective values of 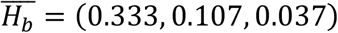 and 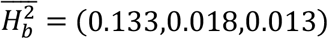. Using the values described above, we estimated the correlation of QTL substitution effects across breeds, 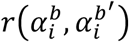, using equation (7).

Similarly, the correlation of QTL substitution effects a certain number of generations away, 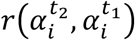, was obtained applying Equation (9). An increase Δ*F* = 0.01 per generation was assumed, for the same assumed variances and allele frequency distributions.

#### Estimates of correlation across populations using literature values

The results for reasonable guesses of the distribution of allele frequencies at QTL are in Table 4. The correlation of QTL substitution effects across breeds was roughly 0.85 across both species (pigs and cattle) and all assumed distributions of gene frequencies. These numbers are higher than scarce estimates of genetic correlations in the literature: 0.4 to 0.8 for production traits (Karoui *et al*. 2012), 0.3 to 0.5 for milk yield (Legarra *et al*. 2014) and 0.6 for body condition (Porto-Neto *et al*. 2015), and we consider that this is because across-breeds genetic correlations also include genotype by environment interactions. It may be argued that these studies used SNPs instead of true QTLs; however, Wientjes et al. (2017) in a simulation study obtained unbiased estimates of correlation across QTLs using SNP markers for a true correlation of 0.5, and (Cuyabano *et al*. 2018) showed analytically that when SNP markers yield correct estimates of true relationships at QTL, estimates of genetic parameters are unbiased.

**Table 4:** Estimates of correlations of QTL effects across breeds based on values from Table 3 and Equation 7, for different distributions of QTL frequencies

| Species | Uniform | Hill<sup>1</sup> | Extreme<sup>2</sup> |
| --- | --- | --- | --- |
| Cattle | 0.83 | 0.82 | 0.84 |
| Pigs | 0.87 | 0.85 | 0.88 |

<sup>1</sup> U-shaped distribution with effective population size of 50
<sup>2</sup> Beta(0.04, 0.04) distribution

Concerning decrease of correlations across generations, results (Table 5) show a slow decrease, and this decrease is, again, not sensitive to the assumed distribution of allelic frequencies. Values in Table 5, quite close to 1, somehow disagree with few existing estimates of across-generation genetic correlations, clearly lower than 1 (Tsuruta *et al*. 2004; Haile-Mariam and Pryce 2015). Again, part of the literature estimates of correlation much lower than 1 is probably due to non-additive gene action, and part due to genotype by environment interactions.

**Table 5:** Correlation of QTL effects within breed across time, based on values from Table 3 and Equation 7, for different distributions of QTL frequencies

| Species | Distance in generations | Uniform | Hill<sup>1</sup> | Extreme<sup>2</sup> |
| --- | --- | --- | --- | --- |
| Cattle | 1 | 0.98 | 0.97 | 0.98 |
|  | 2 | 0.97 | 0.94 | 0.97 |
|  | 5 | 0.93 | 0.87 | 0.93 |
|  | 10 | 0.86 | 0.78 | 0.87 |
| Pigs | 1 | 1.00 | 0.99 | 1.00 |
|  | 2 | 0.99 | 0.98 | 0.99 |
|  | 5 | 0.98 | 0.96 | 0.97 |
|  | 10 | 0.96 | 0.93 | 0.93 |

<sup>1</sup> U-shaped distribution with effective population size of 50
<sup>2</sup> Beta(0.04, 0.04) distribution

## DISCUSSION

The fact that most genetic variation is statistically additive is well known. However, this does not imply equality of statistical additive effects across populations, something that is possibly reflected in the difficulty of predicting breeding values or disease risks across (distinct) populations, either in animal or human genetics. Only recently has non-additive biological gene action been identified as playing a role in determining the degree to which prediction across populations is successful (Dai *et al*. 2020; Duenk *et al*. 2020). These authors presented result of simulations, where the problem is that the results are scenario specific and do not allow a good appraisal of the different factors. Having a theory allows to derive more general expressions and understanding factors influencing across-population differences in substitution effects.

In this work, we present a general, formal framework that does not depend on specific hypothesis regarding gene action or order of the epistatic interactions. In our derivation, we approached *high-order functional* epistasis by Taylor expansions, leading to expressions that involve only *low-order statistical* additive epistasis (actually pairwise). Including one extra term in our Taylor expansion would include *three-loci statistical* epistasis and so on, but extra terms would lead to more difficult expressions (covariance of allele frequencies across loci and populations), and it is expected that the magnitude of the statistical effects is smaller and smaller with higher orders of interactions (Mäki-Tanila and Hill 2014). We find reasonable agreement with our simulation and also with values from literature. However, our simulations may not be particularly realistic, something that would require considerable thinking on how to simulate biologically meaningful epistasis mechanisms for a variety of traits. We see them as building blocks of non-additive architecture. At any rate, the three scenarios generated sizeable non-additive variation which is a challenging case for our expressions.

Three main factors influence the correlation of substitution effects between populations *r*(*α^b^, α^b′^*): the genetic similarity of the two populations, the magnitudes of additive, dominance and additive by additive variances, and the distribution of allele frequencies at QTL. We consider that showing explicitly these three factors is an achievement, as their role is implicit, yet not explicitly shown, in previous works in simulated and real data (Martin *et al*. 2019; Dai *et al*. 2020; Duenk *et al*. 2020). Now we discuss these three factors.

The distance across populations is summarized by Nei’s minimum genetic distances *D_b,b′_* or *F_st_* indexes. Under pure drift scenarios, these depend on divergence times and effective population sizes (Weir and Hill 2002; Bonhomme *et al*. 2010; Walsh and Lynch 2018).

The factors 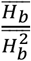 and 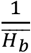 are weighting factors on dominance and additive by additive variances. If the allele frequencies are modelled using symmetric *Beta*(*a, a*) distributions (see Appendix, section 1.11), these become 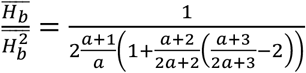 and 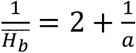. The first is bounded between 3 (for U-shaped distributions) and 2 (for peaked distributions), so that dominance variation does not play a big role in the difference between substitution effects across populations unless dominance variation is much larger that additive variation, something that seems unlikely based on theory and estimates. However, the second weight, due to epistasis, is not bounded and is large for small values of *a* (*e.g*. 27 for Beta(0.04,0.04), and in this case functional epistasis plays a strong role in additive variation for U-shaped distributions (Hill *et al*. 2008). The spectra of allele frequencies of causal mutations is subject to large debate but there seems to be a consensus that low frequency mutations make non-negligible contributions to genetic variance (Eyre-Walker 2010; Gibson 2012). It is unknown if this leads to extreme values of 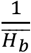.

The particular case of purifying selection deserves some attention. For a mutant recessive deleterious allele with fitness 1 – *s*, the substitution effect is *α* = −*ps* so, in principle, *α* changes across populations – this is similar to our simulated scenario of Complete Dominance. However, if the locus is truly at equilibrium in all populations, then 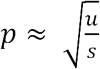 (Crow and Kimura 1970 chap. 6). Provided that *s* and *u* are identical across populations, the allele frequency *p and* the substitution effect *α* should be identical across populations. However, *s* is not necessarily homogeneous across populations, likely depending on the environment.

The previous result (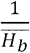 needs to be large for biological dominance to play a role) clarifies the findings of Duenk *et al*. (2020) and Dai *et al*. (2020) that it is mainly biological epistasis that generates changes in additive substitution effects across populations. For instance, when one population drifts, the additive by additive statistical variation enters into additive variation (Hill *et al*. 2006; Walsh and Lynch 2018).

Thus, 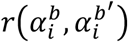 will be very low only if (1) populations are distinct, (2) there is *statistical* additive by additive variation and (3) QTL allele frequency distributions are U- or L-shaped. As for the magnitudes of non-additive variance, in most conceivable cases 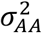 and 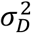 are of magnitudes at most like additive variances, but more often less than half (Fuerst and Sölkner 1994; Palucci *et al*. 2007; Hill *et al*. 2008; Mäki-Tanila and Hill 2014; Vitezica *et al*. 2018), so in practice the most limiting factor is distance across breeds. Indeed, there are very few accurate estimates of non-additive variation although more are becoming available. We argue that then, and based also in our results, it will be possible to make theory and data-based choices on the possibilities of using predictions and GWAS results across populations.

Then, for reasonable assumptions about allele frequency distribution of QTLs, we have shown that the correlation of substitution effects across populations is typically around 0.8 or higher, which is higher than scarce estimates of genetic correlations across populations available in the literature, which range from 0.3 to 0.8 (Karoui *et al*. 2012; Legarra *et al*. 2014; Porto-Neto *et al*. 2015; Xiang *et al*. 2017) the difference being due, probably, to genotype by environment interaction. Our results seem rather robust to different distributions of allele frequencies. These values are high but not 1, which raises the question of how to conceive optimal strategies for across populations predictions (*e.g*. more data within breed or finer locations of causal variants across breed). This is of practical relevance, *e.g*. for genomic predictions in livestock improvement, but also in human genetics *e.g*. for the use of European-based Polygenic Risk Scores in individuals from other ancestries (Martin *et al*. 2019).

Similar results apply to the same populations across generations, in which case the correlation of substitution effects across generations goes from 0.99 (next 1-2 generations) to 0.80 (10 generations). This illustrates that, even if genomic predictions would use QTLs or markers in very tight LD with QTLs, there would still be, in the long run, a need for a continuous system of data collecting and re-estimation of effects.

In the simulations we confirmed that our estimates are reasonably good although not perfect. They are, depending on the scenario and target (random allele or mutant allele), almost unbiased, slightly biased upwards, or biased downwards. There are several reasons for the disagreement. The most obvious one is the inherent approximation of the Taylor series expansion. Secondly, splitting variances such as 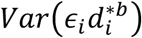 or 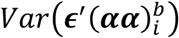 into basic components implies either a strong assumption of multivariate normality and independence or a less strong one of “expectation and variance-independence” (Bohrnstedt and Goldberger 1969). For instance, it is assumed that the variance of 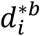 is not related to the magnitude of the difference of allele frequencies *ϵ_i_*, but this is not necessarily true. Third, it is further assumed, in the factorization of genetic variances, that QTL effects and allele frequencies are independent. We have also ignored that the change *ε* is proportional to heterozygosity and therefore small at extremes values of allele frequencies.

Another factor that we did not consider is the empirical evidence of an inverse relationship between heterozygosity and absolute effect at the locus (Park *et al*. 2011). It is unclear how this would affect our findings. A (rather extreme) functional gene action that generates larger *α* at extreme frequencies is overdominance, which is similar to our “dominance” scenario.

As for our estimators of the correlation across “negative” alleles *r_mutant_*(*α^b^, α^b′^*), they are less robust that the estimators for a “random” allele *r_random_*(*α^b^, α^b′^*). The reason for this is that obtaining *r_mutant_*(*α^b^, α^b′^*) from *r_random_*(*α^b^, α^b′^*) involves a further approximation, the normality of *α*.

## CONCLUSIONS

We presented a coherent, approximate theory, that does not invoke any particular mechanism of gene action, to explain and appraise the change in magnitude of (additive) QTL substitution effects across populations and generations. The theory gives good approximate estimates of this correlation, that needs to be otherwise explicitly estimated. More importantly, the theory shows that the main sources for the change of effects are relationships across populations, magnitudes of additive and first-order non-additive variances (dominance and additive by additive), and spectra of allele frequencies. These findings provide better understanding of the properties of genomic prediction methods and of quantitative genetics in general.

## Data Availability Statement

Appendix associated with this manuscript is in https://figshare.com/articles/online_resource/Appendix_to_Genetics_paper/15133947.

Code and files for simulations can also be found in https://figshare.com/articles/software/scripts_zip/14509956. All other results can be reproduced using equations and figures from the text.

## ACKNOWLEDGMENTS

Part of this work was while Y. Wientjes was visiting the GenPhySE unit at INRAE, Toulouse, financed by the Netherlands Organisation of Scientific Research (NWO). Authors thank INRA SelGen metaprograms EpiSel and EpiFun. This project has received funding from the European Unions’ Horizon 2020 Research & Innovation programme under grant agreement N°772787 – SMARTER. We are grateful to the Genotoul Bioinformatics Platform Toulouse Midi-Pyrenees (Bioinfo Genotoul) for providing computing and storage resources. We thank editor and reviewers for very detailed feedback, and MA Toro, A Caballero, S Boitard and M Bonhomme for advice. A.L., Z.G.V. and C.A.G.B. want to dedicate this paper to his friend Eduardo Fernández, who could discuss substitution effects while driving through traffic jams, and who left us too soon.

## Notes

### Competing Interest Statement

The authors have declared no competing interest.

