## Appendix for "The Correlation of Substitution Effects Across Populations and Generations in the Presence of Non-Additive Functional Gene Action" (from another version of this preprint: version 2, 2021-04-30)

* INRAE/INP, UMR 1388 GenPhySE, 31326, Castanet-Tolosan, France.
† Departamento de Producción Animal, Facultad de Agronomía, Universidad de Buenos Aires, C1417DSQ, Buenos Aires, Argentina.
‡ Wageningen University & Research, Animal Breeding and Genomics, 6700 AH Wageningen, The Netherlands.

Here we present detailed derivations of many aspects of this work.

1.1 Taylor series expansion of substitution effects

The function $\alpha_i^{b'} \approx f(\alpha_i^b, \boldsymbol{\epsilon})$, constructed as a Taylor series expansion around $\alpha_i^b$ evaluated at $\boldsymbol{p}^b$, is, e.g. (Walsh & Lynch, 2018, sec. A6.7b):

$$\alpha_i^{b'} \approx \alpha_i^b + \boldsymbol{\nabla}_i'\boldsymbol{\epsilon} + \frac{1}{2}\boldsymbol{\epsilon}'\boldsymbol{\mathcal{H}}_i\boldsymbol{\epsilon} \dots$$

where $\boldsymbol{\nabla}$ and $\boldsymbol{\mathcal{H}}$ are a vector and a matrix containing, respectively, the gradient and Hessian of $\alpha_i$ with respect to $\boldsymbol{p}$ evaluated at $\boldsymbol{p}^b$ and $\boldsymbol{\epsilon} = \boldsymbol{p}^{b'} - \boldsymbol{p}^b$.

There will be different $\boldsymbol{\nabla}_i$ and $\boldsymbol{H}_i$ per each of the $n$ loci. The elements of $\boldsymbol{\nabla}_i$ are:

$$\boldsymbol{\nabla}_i = \begin{pmatrix} \frac{\partial \alpha_i}{\partial p_1}|(p_i = p_i^b) \\ \dots \\ \frac{\partial \alpha_i}{\partial p_i}|(p_i = p_i^b) \\ \dots \\ \frac{\partial \alpha_i}{\partial p_n}|(p_i = p_i^b) \end{pmatrix} = 2\begin{pmatrix} (\alpha\alpha)_{i1}^b \\ \\ -d_i^{*b} \\ \\ (\alpha\alpha)_{in}^b \end{pmatrix}$$

For instance, $(\alpha\alpha)_{i1} = \frac{1}{2}\frac{\partial \alpha_i}{\partial p_1}$ and therefore $\frac{\partial \alpha_i}{\partial p_1}|(p_i = p_i^b) = 2(\alpha\alpha)_{i1}^b$. In other words, the vector of first derivatives of $\alpha_i$ can be written as function of statistical effects in a focal ("*b*") population with allele frequencies $p_i^b$. In the 1st order expansion, these effects are additive by additive $(\alpha\alpha)$ (of a locus with other loci) and dominant $d^*$ (of a locus with itself). The *b* in the notation $(\alpha\alpha)_{i1}^b$ implies that the Taylor series is expanded around a population with $\boldsymbol{p} = \boldsymbol{p}^b$.

The elements of $\boldsymbol{\mathcal{H}}_i$ for a pair of other loci $j$ and $k$ are:

$$\boldsymbol{\mathcal{H}}_{i(jk)} = \left(\frac{\partial \alpha_i}{\partial p_j p_k}|(p_i = p_i^b)\right) = 4(\alpha\alpha\alpha)_{ijk}^b$$

For $i \neq j \neq k$. If $i = j = k$, then $\boldsymbol{\mathcal{H}}_{i(jk)}$ is null and if $j = k$, then $\boldsymbol{\mathcal{H}}_{i(jk)}$ is the additive by dominance epistatic interaction $(-\alpha d)_{ij}$ .

We are going to ignore these higher-order interactions (2nd order including dominance, and 3-way and higher) for mathematical convenience, but also because such statistical interactions are expected to be small (Mäki-Tanila & Hill, 2014). Therefore, from the Taylor series expansion above, we will consider only the first two terms, i.e. $\alpha_i^{b'} \approx \alpha_i^b + \boldsymbol{\nabla}_i'\boldsymbol{\epsilon}$. Written as function of substitution effects this gives, for locus $i$,

$$\alpha_i^{b'} \approx \alpha_i^b + 2\epsilon_i\left(-d_i^{*b}\right) + 2\boldsymbol{\epsilon}'(\boldsymbol{\alpha\alpha})_i^b$$

Where $(\boldsymbol{\alpha\alpha})_i^b$ is a vector containing epistatic substitution effects of locus $i$ with the rest of loci. By convention and to simplify notation we set $(\alpha\alpha)_{ii}^b = 0$ (this corresponds to the interaction of a locus with itself, i.e. the dominance effect $d_i^{*b}$). Note that $\alpha_i^b$, $d_i^{*b}$ and $(\boldsymbol{\alpha\alpha})_i^b$ do *not* depend on allele frequencies in $b'$ (as they are defined for the focal population $b$ with allele frequencies $\boldsymbol{p}^b$).

The expression above also shows, without invoking any explicit "functional" effect, that the lower level statistical effects in a reference framework (or set of allele frequencies) contain higher order statistical effects in another framework (Álvarez-Castro & Carlborg, 2007).

### 1.2 Expansion of $Var\left(\alpha_i^b + 2\epsilon_i\left(-d_i^{*b}\right) + 2\boldsymbol{\epsilon}'(\boldsymbol{\alpha\alpha})_i^b\right)$

First,

$$Var\left(\alpha_i^b + 2\epsilon_i\left(-d_i^{*b}\right) + 2\boldsymbol{\epsilon}'(\boldsymbol{\alpha\alpha})_i^b\right) = Var\left(\alpha_i^b\right) + 4Var\left(\epsilon_i d_i^{*b}\right) + 4Var\left(\boldsymbol{\epsilon}'(\boldsymbol{\alpha\alpha})_i^b\right)$$

as cross-products $Cov\left(\alpha^b, \epsilon_i\left(-d_i^{*b}\right)\right)$ and $Cov\left(\alpha^b, \boldsymbol{\epsilon}'(\boldsymbol{\alpha\alpha})_i^b\right)$ are 0 because statistical effects ($\alpha_i^b$, $d_i^{*b}$ and $(\boldsymbol{\alpha\alpha})_i^b$) are mutually orthogonal. For $Var\left(\epsilon_i d_i^{*b}\right)$, we use (under certain assumptions, roughly speaking of independence of $\epsilon_i$ and $d_i^{*b}$) result [9] in Bohrnstedt and Goldberger (1969) : $Var(xy) = E^2(x)Var(y) + E^2(y)Var(x) + Var(x)Var(y)$, leading to

$$Var\left(\epsilon_i d_i^{*b}\right) = E^2(\epsilon_i)Var\left(d_i^{*b}\right) + E^2\left(d_i^{*b}\right)Var(\epsilon_i) + Var(\epsilon_i)Var\left(d_i^{*b}\right)$$
$$= Var(\epsilon_i)\left(Var\left(d_i^{*b}\right) + E^2\left(d_i^{*b}\right)\right)$$

as $E(\epsilon_i) = 0$ (as changes are assumed to be random and not directional). In turn, $E^2\left(d_i^{*b}\right) = \mu_{d,b}^2$ from above which yields $Var(\epsilon_i)\left(\sigma_{d,b}^2 + \mu_{d,b}^2\right)$.

The expansion of $4Var\left(\boldsymbol{\epsilon}'(\boldsymbol{\alpha\alpha})_i^b\right)$ involves the variance of a quadratic form as shown below (section 1.3).

### 1.3 Variance of a quadratic form

The expression for the variance $Var(\boldsymbol{x}_1'\boldsymbol{x}_2)$ of a bilinear form $(\boldsymbol{x}_1'\boldsymbol{x}_2)$ for two vectors, with joint covariance matrix $Var\begin{pmatrix}\boldsymbol{x}_1\\ \boldsymbol{x}_2\end{pmatrix} = \boldsymbol{C} = \begin{pmatrix}\boldsymbol{C}_{11} & \boldsymbol{C}_{12}\\ \boldsymbol{C}_{21} & \boldsymbol{C}_{22}\end{pmatrix}$ and $E\begin{pmatrix}\boldsymbol{x}_1\\ \boldsymbol{x}_2\end{pmatrix} = \begin{pmatrix}\boldsymbol{0}\\ \boldsymbol{0}\end{pmatrix}$, is equal to $Var(\boldsymbol{x}_1'\boldsymbol{x}_2) = tr(\boldsymbol{C}_{21})^2 + tr(\boldsymbol{C}_{22}\boldsymbol{C}_{11})$ (Searle, 1971, Chapter 2). For our specific case, we are interested in $Var\left(\boldsymbol{\epsilon}'(\boldsymbol{\alpha\alpha})_i^b\right)$ with $E\begin{pmatrix}\boldsymbol{\epsilon}'\\ (\boldsymbol{\alpha\alpha})_i^b\end{pmatrix} = \begin{pmatrix}\boldsymbol{0}\\ \boldsymbol{0}\end{pmatrix}$ and $Var\begin{pmatrix}\boldsymbol{\epsilon}'\\ (\boldsymbol{\alpha\alpha})_i^b\end{pmatrix} = \begin{pmatrix}Var(\boldsymbol{\epsilon}') & \boldsymbol{0}\\ \boldsymbol{0} & Var\left((\boldsymbol{\alpha\alpha})_i^b\right)\end{pmatrix}$ where we assume $Cov\left(\boldsymbol{\epsilon}', (\boldsymbol{\alpha\alpha})_i^b\right) = \boldsymbol{0}$. Thus, the above expression reduces to

$$Var(\boldsymbol{\epsilon}'(\boldsymbol{\alpha\alpha})_i^b) = tr\left(Var((\boldsymbol{\alpha\alpha})_i^b)Var(\boldsymbol{\epsilon}')\right) = tr\left(Var(\boldsymbol{\epsilon}')Var((\boldsymbol{\alpha\alpha})_i^b)\right)$$

1.4 Variance due to dominance effects and inbreeding depression

In a HWE population with dominance deviations, the genetic variance due to dominance deviations is

$$\sigma_D^2 = 4\sum p_i^2 q_i^2 (d_i^*)^2$$

To account for directional dominance, consider $E(d_i^*) = \frac{1}{n}\sum(d_i^*) = \mu_d$, $Var(d_i^*) = \sigma_d^2$. Then we can define "centered" $d_i^{*(c)} = d_i^* - \mu_d$, with $Var\left(d_i^{*(c)}\right) = \sigma_d^2$. If there are $n$ loci, then the effect of inbreeding per unit of homozygosity is $b = -\mu_d n$ . Now we can decompose $\sigma_D^2 = 4\sum p_i^2 q_i^2 (d_i^*)^2 = 4\sum p_i^2 q_i^2 \left(d_i^{*(c)}\right)^2 - 4\sum p_i^2 q_i^2 (2d_i^{*c}\mu_D) + \mu_d^2 4\sum p_i^2 q_i^2 = 4\sum p_i^2 q_i^2 \left(d_i^{*(c)}\right)^2 + \mu_d^2 4\sum p_i^2 q_i^2$ because $4\sum p_i^2 q_i^2 (2d_i^{*c}\mu_D) = 0$ across loci. Assuming independence of $p_i^2 q_i^2$ and $d_i^{*(c)}$, then $\sigma_D^2 = 4\sum p_i^2 q_i^2 (\sigma_d^2 + \mu_d^2)$ or in other terms, $\sigma_D^2 = n\overline{H^2}(\sigma_d^2 + \mu_d^2)$ where $\overline{H^2}$ is average squared heterozygosity.

1.5 Variance of statistical effects

We use the fact that the population variances are function of the variances of the different effects (Maki-Tanila and Hill, 2014) and moments of heterozygosities. First, we define functions of heterozygosities for all $n$ loci at the focal population. The average heterozygosity across loci is:

$$\overline{H_b} = \frac{1}{n}\sum 2p_i^b(1 - p_i^b)$$

The average squared heterozygosity is

$$\overline{H_b^2} = \frac{1}{n}\sum \left(2p_i^b(1 - p_i^b)\right)^2$$

and the average cross-product of heterozygosities across all pairs of distinct loci is

$$\overline{HH_b} = \frac{1}{0.5n(n-1)}\sum_i \sum_{j>i} 2p_i^b(1 - p_i^b)2p_j^b(1 - p_j^b)$$

In fact $n^2(\overline{H_b})^2 = 2n(n-1)\overline{HH_b} + n\overline{H_b^2}$ and from here we make the approximation $\overline{HH_b} \approx \frac{1}{2}\overline{H_b H_b}$.

If we knew these functions of heterozygosities, we could obtain, from estimates of additive, dominant and additive by additive variances, the variance of statistical additive, dominant and additive by additive effects:

$$Var(\alpha_i^b) = \sigma_{\alpha,b}^2 = \frac{\sigma_A^2}{n\overline{H_b}}$$

$$Var(d_i^{*b}) + E^2(d_i^{*b}) = \sigma_d^2 + \mu_d^2 = \frac{\sigma_D^2}{n\overline{H_b^2}}$$

$$Var((\alpha\alpha)_{i,j,j>i}^b) = \sigma_{(\alpha\alpha,b)}^2 = \frac{\sigma_{AA}^2}{n(n-1)\overline{HH_b}} \approx 2\frac{\sigma_{AA}^2}{n^2\overline{H_b}\,\overline{H_b}}$$

$$Var((\boldsymbol{\alpha\alpha})_i^0) = \boldsymbol{I} \otimes \sigma_{(\alpha\alpha,b)}^2 \approx 2\boldsymbol{I}\frac{\sigma_{AA}^2}{n^2\overline{H_b}\,\overline{H_b}}$$

All variances and effects refer to the focal population with allele frequencies $p^b$ and effects $\alpha^b$. Note that we assume HWE and LE.

1.6 Nei's minimal genetic distance $D_{b,b'}$ as function of $F_{ST}$ and $\bar{H}_b$

The objective is to put $D_{b,b'}$ as a function of assumed known parameters $F_{ST}$ and $\bar{H}_b$ (heterozygosity of population $b$). In fact, $F_{ST} = 1 - \frac{0.5(\bar{H}_b + \bar{H}_{b'})}{\bar{H}_a} = \frac{D_{b,b'}}{\bar{H}_a}$ where $\bar{H}_a = \frac{1}{n}\sum_i \left(p_i^{b'}(1-p_i^b) + p_i^b(1-p_i^{b'})\right)$. Assuming $\bar{H}_b \approx \bar{H}_{b'}$ and after some manipulation, this yields $D_{b,b'} = \frac{F_{ST}}{1-F_{ST}}\bar{H}_b$ . Note that although Nei's $F_{ST}$ and Hudson et al. (1992) definitions of $F_{ST}$ differ (Bhatia et al., 2013), their numerator is identical, and identical to $D_{b,b'}$ by definition.

1.7 Correlation of absolute values of bivariate normal

Consider $x$ and $y$ that are multivariate normal with 0 expectation and $Var\begin{pmatrix} x \\ y \end{pmatrix} = \begin{pmatrix} \sigma_x^2 & \sigma_{xy} \\ \sigma_{xy} & \sigma_y^2 \end{pmatrix}$

Now we define the transformed $|x|$ and $|y|$. Using Kan and Robotti (2017) one gets expressions for $E(|x|)$, $Var(|x|)$, and also for $|y|$, and $Cov(|x|,|y|)$. From here the values of the correlation can be obtained through a rather long expression.

This has been conveniently programmed in R package MomTrunc (Galarza et al., 2020). The r code to obtain the correlation of absolute values from the regular correlation is function `r2rabs` as follows:

```
require("MomTrunc")
foo=function(x)
 cov2cor(meanvarFMD(c(0,0),Sigma=matrix(c(1,x,x,1),2),dist='normal')$varcov)[1,2]
r2rabs = function(r){
 out=c()
 for (i in r){
     out=c(out,foo(i))
 }
 out
}
```

Which translates into the following quadratic:

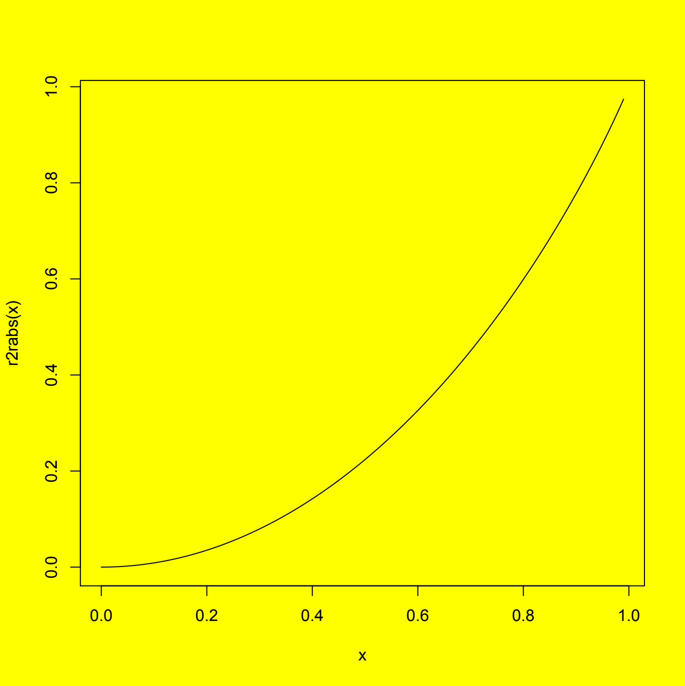


1.8 Simulation

The basic macs command is:

```
macs 400 3e5 -i 100 -t 0.0012 -r 0.001 -I 2 200 200 -ej $t1Over4Ne
2 1 -eN 1.67 10
```

Which translates as: generate 400 sequences of 3e5 bases, 100 times (i.e. 100 DNA stretches). -t specifies the nucleotide diversity per base pair, -r the scaled recombination rate (in 4Ne units), = $\theta = 4Ne\, r$ where $Ne = 300$, the population split into two populations of 200 gametes each (`-I 2 200 200`) at time `$t1Over4Ne` i.e. $t/4Ne$ , -eN specifies that the population had a 10-fold population bottleneck 1.67 generations ago in $4Ne$ units.

This results in the following histogram of allele frequencies for population 1:

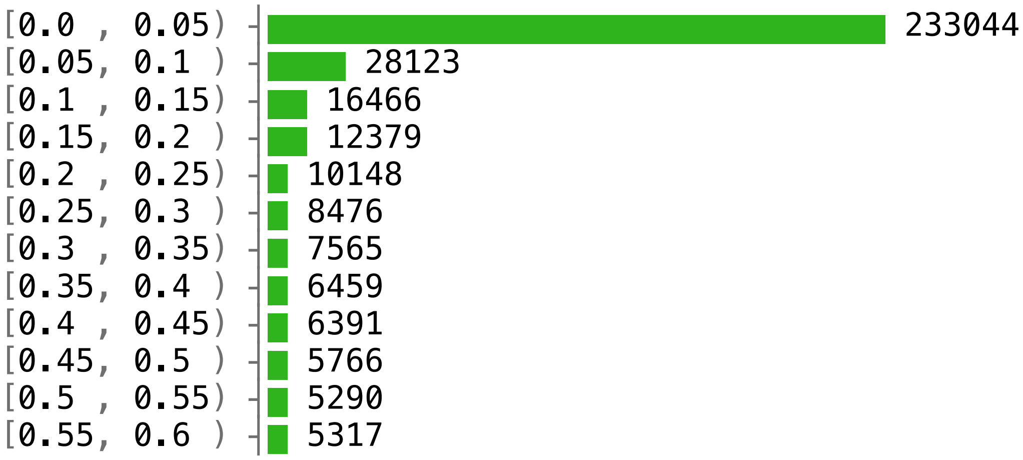

```
[0.6 , 0.65) -█ 4378
[0.65, 0.7 ) -█ 4375
[0.7 , 0.75) -█ 4458
[0.75, 0.8 ) -█ 4200
[0.8 , 0.85) -█ 3792
[0.85, 0.9 ) -█ 3667
[0.9 , 0.95) - 3620
[0.95, 1.0 ) - 3474
[1.0 , 1.05) - 3299
                                        ┘
                    Frequency
```

1.9 <u>Derivation of substitution effects in dominance and epistatic systems</u>

Generally, we use gene content (z-score) coded as {-1,0,1} for the three different genotypes.

*Complete dominance.* In this case we have:

| | cc | Cc | CC |
|---|---|---|---|
| z | $-1$ | 0 | 1 |
| | $a$ | $a$ | 0 |

Here the mean is $a(1-p^2)$ and $\alpha = \frac{1}{2}\frac{\partial E(y)}{\partial p_i} = -ap$ where $p = frequency(C)$. The dominance deviation is $d_i^* = -\frac{1}{2}\frac{\partial \alpha_i}{\partial p_i} = -a/2$.

*Complementary Epistasis.* We can write the 2-locus complementary epistasis model as follows:

| | | cc | Cc | CC |
|---|---|---|---|---|
| | z | $-1$ | 0 | 1 |
| bb | -1 | $a$ | $a$ | 0 |
| bB | 0 | $a$ | $a$ | 0 |
| BB | 1 | 0 | 0 | 0 |

The $n$-locus case generalizes to:

- $a$ if genotypes at all $n$ loci are NOT "upper case upper case"
- 0 otherwise

in other words

$$y = [z_1 \neq 1][z_2 \neq 1] \dots [z_n \neq 1]a$$

where $z$ is the genotype, the operator "[ ]" is Iverson brackets notation (1 if true and 0 otherwise). Then, assuming independence across loci (i.e. LE), we generalize to $n$ loci:

$$E(y) = E([z_1 \neq 1][z_2 \neq 1][z_3 \neq 1]a)$$
$$= aE([z_1 \neq 1])E([z_2 \neq 1])E([z_3 \neq 1]) \dots E([z_n \neq 1])$$

Now we obtain $E([z_1 \neq 1])$. The expression $[z_1 \neq 1]$ can be written as $[z_1 \neq 1] = 1 - \frac{(z_1+z_1^2)}{2}$ (this is not really needed). Alternatively, the expectation $E([z_1 \neq 1]) = q_1^2 + 2p_1q_1 = 1 - p_1^2$ is obtained looking at the following table:

| | z-score | $[z_1 \neq 1]$ | frequency | sum |
|---|---|---|---|---|
| bb | -1 | 1 | $q^2$ | $q^2$ |
| bB | 0 | 1 | $2pq$ | $2pq$ |
| BB | 1 | 0 | $p^2$ | 0 |

Thus, $E(y) = a(1-p_1^2)(1-p_2^2)(1-p_3^2) \dots (1-p_n^2)$

Then we can get the different $\alpha$ as *e.g.* for the first locus we have

$$\alpha_1 = \frac{1}{2}\frac{\partial E(y)}{\partial p_1} = -ap_1\Pi_{j>1}\left(1-p_j^2\right)$$

If we call $K = \Pi\left(1-p_j^2\right)$ then

$$\alpha_1 = \frac{1}{2}\frac{\partial E(y)}{\partial p_1} = -a\frac{p_1}{(1-p_1^2)}K$$

and generally

$$\alpha_i = \frac{1}{2}\frac{\partial E(y)}{\partial p_i} = -ap_i\Pi_{j\neq i}\left(1-p_j^2\right) = -a\frac{p_i}{(1-p_i^2)}K$$

The dominance deviation is

$$d_i^* = -\frac{1}{2}\frac{\partial \alpha_i}{\partial p_i} = \frac{a}{2}\Pi_{j\neq i}\left(1-p_j^2\right) = \frac{a}{2}\frac{1}{(1-p_i^2)}K$$

The $i$ – by – $j$ epistatic effect $(\alpha\alpha)_{ij}$ is

$$(\alpha\alpha)_{ij} = \frac{1}{2}\frac{\partial \alpha_i}{\partial p_j} = \frac{1}{2}\frac{\partial}{\partial p_j}\left(-ap_i\left(1-p_j^2\right)\Pi_{k\neq i,j}(1-p_k^2)\right) =$$

$$ap_ip_j\Pi_{k\neq i,j}(1-p_k^2) = a\frac{p_ip_j}{(1-p_i^2)\left(1-p_j^2\right)}K$$

Note that the expression is oriented: $p_i$ is the frequency of the "recessive" allele (say B) at locus $i$.

*Multiplicative*. Consider, for instance, the following values of the genotypic value for two loci with their respective frequencies:

| | | cc | Cc | CC |
|---|---|---|---|---|
| | z | $-1$ | 0 | 1 |
| bb | -1 | $a$ | 0 | $-a$ |
| bB | 0 | 0 | 0 | 0 |
| BB | 1 | $-a$ | 0 | $a$ |

The genotypic value can be expressed as the product of z values at each loci, i.e. $G = z_1 z_2$ for two loci. The generalization to several loci is immediate as $\prod_{i=1,n} z_i$ . The average genotypic value considering *n* loci is, assuming Cov($Z_i, Z_j$) = 0 (LE):

$$\mu = E(Z_1 Z_2 Z_3 \dots Z_n) = E(Z_1)\,E(Z_2)\,E(Z_3) \dots E(Z_n) \;\; [1]$$

In general, $E(Z_i) = p_i^2 - q_i^2 = p_i - q_i = 2p_i - 1$ and consequently,

$$\mu = \prod_{i=1}^{n} (2p_i - 1)$$

Following Kojima's definition, the additive substitution effect at locus *i* is the first derivative of $\mu$ :

$$\alpha_i = \frac{1}{2}\frac{\partial \mu}{\partial p_i} = \frac{1}{2}\frac{\partial (2p_i - 1)}{\partial p_i}\, \Pi_{i \neq j}\left(2p_j - 1\right) = \prod_{j \neq i}\left(2p_j - 1\right)$$

Or, calling $K = \Pi\left(2p_j - 1\right)$

$$\alpha_i = \frac{1}{2p_i - 1} K$$

The dominance deviation is 0 as expected:

$$d_i^* = -\frac{1}{2}\frac{\partial \alpha_i}{\partial p_i} = -\frac{1}{2}\frac{\partial}{\partial p_i}\prod_{j \neq i}\left(2p_j - 1\right) = 0$$

The additive by additive effect is:

$$(\alpha\alpha)_{ij} = \frac{1}{2}\frac{\partial \alpha_i}{\partial p_j} = \prod_{k \neq i,j} (2p_k - 1) = \frac{1}{(2p_i - 1)\left(2p_j - 1\right)} K$$

1.10 <u>Derivation of the moments of functions of allele frequencies assuming beta distributions</u>

We need the moments

$$\bar{H} = E\left(2p_i(1 - p_i)\right)$$
$$\overline{H^2} = E\left(4p_i(1 - p_i)p_i(1 - p_i)\right)$$

and

$$\overline{HH} = E_{i>j}\left(2p_i(1 - p_i)2p_j\left(1 - p_j\right)\right)$$

from the parameters of the $Beta(\alpha, \beta)$ distribution of the allele frequencies $p$. The moments can be obtained using the Moment generating function of the Beta distribution such that $(p^k) = \prod_{r=1}^{k} \frac{\alpha + r - 1}{\alpha + \beta + r - 1}$ . For instance

$$\bar{H} = 2E\left(p(1 - p)\right) = 2E(p) - 2E(p^2) = 2\frac{\alpha\beta}{(\alpha + \beta)(\alpha + \beta + 1)}$$

Which for $\alpha = \beta = a$ yields

$$\overline{H} = \frac{2a^2}{2a(2a+1)} = \frac{a}{(2a+1)}$$

then

$$\overline{H^2} = E\big(4p(1-p)p(1-p)\big) = 4E(p^2 - 2p^3 + p^4)$$
$$= 4\frac{\alpha}{\alpha+\beta}\frac{\alpha+1}{\alpha+\beta+1}\left(1 + \frac{\alpha+2}{\alpha+\beta+2}\left(\frac{\alpha+3}{\alpha+\beta+3} - 2\right)\right)$$

Which for $\alpha = \beta = a$ yields

$$\overline{H^2} = 2\frac{a+1}{2a+1}\left(1 + \frac{a+2}{2a+2}\left(\frac{a+3}{2a+3} - 2\right)\right)$$

Then, is $\overline{HH} = E_{i>j}\left(2p_i(1-p_i)2p_j(1-p_j)\right)$, that we approximated as $\overline{HH} \approx \frac{1}{2}(\overline{H})^2 = \frac{1}{2}\left(\frac{a}{(2a+1)}\right)^2$

Finally,

$$\frac{\overline{H^2}}{\overline{H}} = 2\frac{a+1}{a}\left(1 + \frac{a+2}{2a+2}\left(\frac{a+3}{2a+3} - 2\right)\right)$$
